# ARIH1 deficiency impairs spatial learning and memory via GIRK2 upregulation in hippocampal CaMKII-expressing neurons in mice

**DOI:** 10.1101/2025.03.10.625121

**Authors:** Chunsheng Zhang, Zhiruo Sun, Xiaowen Chen, Huicui Yang, Jiaojiao Chen, Zhong Ding, Xuechu Zhen

## Abstract

Alterations of Ariadne RBR E3 Ubiquitin Protein Ligase 1 (ARIH1), a human homologue of *Drosophila* Ari, have been associated with a number of human diseases. Given the importance of ubiquitin-proteasome system in learning and memory, whether ARIH1 involves in the process has not been explored. Here we report that ARIH1-deficent mice exhibited a defect in learning and memory evidenced in Morris water maze and in novel object recognition tests without changes in basal motor activity, anxiety, and depressive behaviors. We found that ARIH1 deficiency resulted in an upregulation of G protein-gated inwardly rectifying potassium channel 2 (GIRK2) in dorsal hippocampus that was attributed to the impaired ubiquitination and degradation. Locally injection of ARIH1-expressing lentivirus to restore the ARIH1 expression of dorsal hippocampus in ARIH1^+/-^ mice restored the impaired learning and memory. Moreover, selective knockdown ARIH1 in dorsal hippocampal calcium-calmodulin-dependent protein kinase II (CaMKII)-expressing neurons, but not for parvalbumin^+^ (PV) or somatostatin^+^ (SST) neurons, in naïve mice was sufficient to mimic the damage in learning and memory of ARIH1^+/-^ mice. Lastly, we demonstrated that systemically or locally inhibition of GIRK activity was able to improve ARIH1 deficiency-induced decline of learning and memory in ARIH1^+/-^ mice. The present study discovered the clear role of ARIH1 in mediating learning and memory, defect of ARIH1 resulted in upregulation of GIRK2 in hippocampal CaMKII-expressing neurons via modulating the ubiquitination and degradation GIRK2.

## Introduction

The E3 ubiquitin ligase ARIH1 (also known as Human Homolog of Ariadne-1, HHARI) was firstly identified in *Drosophila* as gene Ari-1 (Ariadne-1) [1]. Recent studies have revealed the important functional roles of ARIH1 in against genotoxic stress, parkin-independent mitophagy, anti-viral effect, as well as promoting cancer progression or anti-tumor immunity [2–8]. In humans, variants in ARIH1 are associated with the developing aortic aneurysms [9], and ARIH1 mutation is observed in patients with leukemia [10].

Study from *Drosophila* has shown that survivors of Ari-1 null alleles exhibited the impaired neuronal development and motor functions [11]. Loss-of-function mutant of Ari-1 in flies resulted in a low spontaneous neurotransmitter release through its substrate N-ethylmaleimide sensitive factor (NSF) [12]. Interestingly, postmortem brain examination revealed ARIH1 as a component of Lewy bodies, a pathological hallmark of some neurodegenerative disorders especially for Parkinson’s disease [13, 14]. ARIH1 was further shown to mediate the formation of aggresomes [13]. Furthermore, elevated ARIH1 mRNA was reported in the hippocampus of female rats after chronic alcohol administration [15]. All these data revealed that ARIH1 may be an important regulator in brain function.

The ubiquitin–proteasome system (UPS) is composed of the proteasome complex and three classes of enzymes including E1 (ubiquitin-activating), E2 (ubiquitin-conjugating) and E3 (ubiquitin-ligases) enzymes [16]. The UPS is an essential regulator in cellular physiological homeostasis, whilst alteration in UPS contributes to numerous pathological conditions. The importance of UPS in the modulation of learning and memory has been documented via both non-proteolytic and proteasome-dependent proteolytic pathways [17, 18]. In this regard, the role of E3 enzymes in AMPA receptors (AMPAR) regulation has been well-documented in synaptic plasticity and learning and memory [16, 19]. For example, eliminating the expression of E3 ligases RNF167 disrupts the ubiquitination and neuronal membrane expression of AMPAR [20]. Application of proteasome inhibitors was shown to interrupt the hippocampal synaptic plasticity via proteasome-dependent proteolytic mechanism [21, 22]. Mice with E3 ubiquitin ligase RNF220 deficiency exhibit an elevated AMPA receptor (AMPAR) expression, resulting in the impaired synaptic plasticity and learning and memory [23]. Meanwhile, effort to depict the mechanism underlying the E3-regulated learning and memory has been reported. For instance, E3s-regulated expression of Arc (Activity-regulated cytoskeleton-associated protein) was shown to alter AMPAR trafficking [24–26]. The E3-regulated expression of hippocampal histone deacetylase 7 (HDAC7) was shown to associate with the E3-modulating memory formation [27]. Regardless, the molecular details for E3 ubiquitin ligases-regulated learning and memory remain uncharted.

Here, we report that ARIH1-deficient mice exhibited the impaired spatial learning and memory and recognition ability. We further revealed that upregulation of GIRK2 in hippocampal CaMKII-expressing neurons is likely to be responsible for the learning and memory defect in ARIH1-deficient mice.

## Materials and Methods

### Cell lines and animals

HT-22 mouse hippocampus cells stably expressing the Lenti-shRNA (ARIH1-948 and -949) were constructed and maintained in DMEM (Gibco, Thermo-Fisher) supplemented with 5% fetal clone I serum (Hyclone, USA), 1% Antibiotic-Antimycoctic (Life Technologies), and 1 μg/ml puromycin (Sigma). All cell lines were cultured at 37°C with 5% CO_2_.

ARIH1^+/-^ and ARIH1-flox mice were offered by CAM-SU Genomic Resource Center (Soochow University), Pvalb-2A-Cre (NM-KI-200098) and Sst-IRES-Cre (NM-KI-190091) mice were purchased from SMOC (Shanghai Model Organisms Center), and CaMK2α-cre mice was purchased from Cyagen (Suchou, China). ARIH1-PV-cKO and ARIH1-SST-cKO mice were generated by crossing ARIH1-flox mice with Pvalb-2A-Cre and Sst-IRES-Cre, respectively. All mice were bred in SPF laboratory animal facility, and group-housed five/cage under a 12-h light-dark cycle (light on from 6 AM to 6 PM) in stable conditions with food and water ad libitum.

All animal studies and experimental procedures were approved by the Animal Care and Use Committee of Soochow University and were following Guidelines for the Care and Use of Laboratory Animals (Chinese National Research Council, 2006, ethical approval number: 202208A0688) and the “ARRIVE” (Animals in Research: Reporting In Vivo Experiments) guidelines.

### Plasmids, Viruses, and transfections

The Flag-ARIH1 (HG20139-CF) and HA-GIRK2 (HG18578-CY) plasmids were purchased from Sino Biological. The Lenti-ARIH1-EGFP was purchased from OBiO Technology. The Lenti-shRNA (ARIH1-948 and -949)-EGFP and AAV-DIO-shRNA (ARIH1-948)-EGFP were purchased from Genechem (Shanghai, China). Serotype 9 AAV vector was used in this study and all the plasmids were designed and constructed by standard methods.

Briefly, the plasmids and Lipofectamine 2000 (Thermo Fisher Scientific) were respectively diluted in DMEM to desired concentrations. After 5 min incubation, DNA and Lipofectamine solutions were mixed, followed by incubation at room temperature for 30 min, and added to the cells dropwise. Cells were transfected for approximately 48 h prior to being used for the co-IP/immunoblot assays.

Similarly, the Lenti-shRNA (ARIH1-948 and -949)-EGFP and Lenti-NC were transfected into HT-22 cells according to the manufacturer’s protocol for 48 h, before adding puromycin (2.5 μg/ml) in the cell culture media for selection. After 14 days selection, NC-HT-22, 948-HT-22 and 949-HT-22 cells were maintained in cell culture media supplemented with 1 μg/ml puromycin.

### Antibodies

The following antibodies were used: anti-ARIH1 (Ab3891, Abcam), anti-ARIH1 (NBP2-57888, Novus Biologicals), anti-GIRK2 (21647-1-AP, Proteintech), anti-

GAPDH (60004-1-lg, Proteintech), anti-β-actin (66009-1-Ig, Proteintech), anti-Flag (20543-1-AP, Proteintech), anti-HA (51064-2-AP, Proteintech), anti-CaMKII (ab52476, Abcam), anti-GFAP (ab7260, Abcam), anti-Ib1 (011-27991, Wako), anti-Parvalbumin (ab32895, Abcam), and anti-Somatostatin (ab140665, Abcam).

### RT-qPCR

Total RNA was extracted from cells or mouse tissues using RNAiso Plus (TaKaRa, Japan) according to the manufacturer’s instructions. Then, 1 μg RNA was reverse transcribed into cDNA using RT kit (TaKaRa) before cDNA was amplified using specific primers. A StepOnePlus Real-Time PCR System (Applied Biosystems, USA) was used to quantify mRNA expression using SYBR Premix II (TaKaRa, Japan).

The parameters for RT-qPCR were 30s at 95°C, 5s at 95°C, and 30s at 60°C for 40 cycles. GAPDH was used as a reference gene.

### Immunoprecipitation and western blot

The cell extracts or brain tissue homogenates were prepared in lysis buffer (50 mM HEPES, [pH 7.4], 150 mM NaCl, 5 mM EDTA,0.1%Triton, X-100) containing protease inhibitor cocktail (Sigma). Then, 15∼25μg proteins were loaded and separated by SDS-PAGE gel (6%∼12%) electrophoresis before transferring onto a PVDF membrane and probing by specific antibodies. Alternatively, 150∼200μg of lysate were incubated with 1.5μg of antibodies overnight at 4°C, followed by incubation with Protein A/G Magnetic Beads (Selleck) for 4 h at room temperature. The precipitates were washed 3 times with a washing buffer (50 mM Tris, 150 mM NaCl, 0.1% Triton, X-100, pH 7.5), before the immune-precipitated proteins were subjected to SDS-PAGE gel electrophoresis and probed by specific antibodies.

### Stereotaxic surgery and histology

Mice were anesthetized with ketamine (100 mg/kg of body weight) and xylazine (8 mg/kg) by *i.p.* injection and placed in a stereotactic frame. Then, 0.3μl/side of AAV virus (approximately 10^12^ infection units per ml) or 0.6μl/side of lentivirus (approximately 10^8^ infection units per ml) were injected into mouse dorsal hippocampus (coordinates from bregma: 2.18 mm AP, ±2.18 mm ML, −3.2mm DV) using glass microelectrodes at a rate of 30 nl/min. The microelectrode was slowly withdrawn 5 min after the virus infusion. Behavioral experiments were performed 3 weeks after surgery.

Similarly, mice were implanted with canular at the same coordinates, and were subjected to behavioral tests after 1 week recovery.

Brain slices from mice with stereotaxic surgery were counterstained with Hoechst and directly examined under fluorescent microscope. Only data obtained from animals with correct injections were included.

### Behavioral tests

For all the behavioral assays below, mice were allowed to accommodate to the experimental environment for at least 30min before the tests.

#### Open filed

Mice were placed into the arena (40 height x 40 width x 40cm depth) and videotaped for 30min. The total distance (m) and time spend in center (s) of the mice were analyzed using video-tracking software (ANY-maze, USA).

#### Rota Rod

In the training section, mice were place on the instrument for 5min daily with a constant speed of 20rpm for 2 consecutive days. On the third day, mice were placed on the same instrument but with a gradual increasing speed of 4–40rpm, and the latency to fall (s) of mice was recorded.

#### Pole test

A rough-surface wooden pole (60cm height, 1cm diameter) with a ball (2.5cm diameter) on the top was employed in the current study. During the test, the mouse was placed on the ball facing upwards, and the time taken by the mouse to reach the floor with its forelimbs was recorded as the pole climb downtime

(s). Final results are presented as average value of 3 consecutive trails for each mouse.

#### Elevated plus maze

A cross-shaped instrument with two open arms (30 length x 5 cm width) across from each other and perpendicular to two closed arms (30 length x 5 width x 10 cm height) with a center platform (5 length x 5 cm width) was employed in the current study. Briefly, mice are placed on the center platform facing the open arm and allowed to move freely in the maze for 10 min. Mice behavior was videotaped from the top of the instrument, and the number of entries (an entry is defined as the center of mass of the mouse enters the arm) into open arms (counts) and the time spent in open arms (s) are analyzed using video-tracking software (ANY-maze, USA).

#### Tail suspension test

Homemade tail suspension boxes with the dimensions (50 height X 50 width X 15 cm depth) and hooks on the top of box were employed in the current study. Briefly, tapes with same lengths (15-17cm) were used for suspension of the mouse with one end attaching to the tail and the other end attaching to the hook. Mice behavior was videotaped for 6min, and the immobility time (s) of each mouse was analyzed using video-tracking software (ANY-maze, USA).

#### Morris water maze

A circular pool with a diameter of 150cm and a depth of 50cm was employed in the current study. The pool that loaded with opaque water was divided into 4 quadrants with 4 different cues (cards with different shape and color) attached on each wall, and a hidden platform (1.5cm beneath the water surface) was set in the middle of one of the 4 quadrants. A video recording and analyzing system was set above the pool.

On the day for adaptive (day 1), the mouse was placed into the water facing the wall for only once (in the quadrant that opposite to the one set with platform), and the latency to find to platform (escape latency, s) was recorded. If the mouse does not find the platform, the latency was recorded as 60s. In the training section (day 2-7), the mouse was subjected to 4 trails a day (respectively in 4 different quadrants), and each escape latency was recorded. If the mouse finds the platform in 60s, allow the mouse to stay on the platform for another 5s before putting it back to the home cage. If the mouse does not find the platform in 60s, place the mouse on the platform for 20s before returning it back to the home cage. Final results are presented as average value of 4 trails for each mouse a day. On the test day (day 8), the escape latency was recorded after putting the mouse into the water (in the quadrant that opposite to the one set with platform) for the first time, and the times across the platform area (counts) was measured 12hours later, after removing the platform and putting the mouse into the water similarly as the first test.

#### Novel object recognition

The chamber (40 height x 25 width x 25cm depth) with a video recording and analyzing system setting above was employed in the current study. In training section, after exploring the chamber for 5min, allowing the mouse to freely access to two same objects that located near to two corners of the chamber for another 5min before returning the mouse to the home cage. After 30min, one object was replaced by a novel one, and letting the mouse to freely access to both objects for another 5min. The recognition time (s) for each object was analyzed by video-tracking software. Discrimination index was calculated as: recognition time for novel object / (recognition time for novel object + old object).

### Statistical analysis

All data are presented as mean ± SEM. Comparisons between 2 groups were done using unpaired Student’s t test. Comparisons among multiple groups under 1 condition were done using one-way ANOVA and comparisons among multiple groups under 2 different conditions were performed using two-way ANOVA. The Tukey’s and Bonferroni’s multiple-comparison test were used as indicated in the legends for post hoc analyses. Fiji software was used to quantify the fluorescence intensity of IHC images.

## Results

### Deficiency of ARIH1 in mice exhibited an impaired learning and memory

Since the lethality of completely ARIH1 gene knockout (homozygotes, ARIH1^-/-^) [11], we bred heterozygotes mice (ARIH1^+/-^) to test the effect of ARIH1 deficiency on learning and memory. The defect of ARIH1 were confirmed both in qPCR and WB. As expected, both mRNA and protein expression in ARIH1^+/-^ mice were decreased about 50% comparing with that of their WT controls (Figure 1A-1C). We employed two different behavioral paradigms Morris water maze (MWM) (Figure 1D and 1E) and novel object recognition (NOR) test (Figure 1I) to detect the spatial learning and memory and cognitive ability in ARIH1^+/-^ mice. As shown in Figure 1F and 1G, the latency of male ARIH1^+/-^ mice in finding the platform was significantly higher than that of the WT controls. In consistent, the times crossing the platform area for male ARIH1^+/-^ mice was significantly shorter than that of the WT mice (Figure 1H). We found that the impairments were independent of sex, since female ARIH1^+/-^ mice also exhibited a similar deficit in MWM test (SI Figure 1A-1C). In NOR test, we observed that ARIH1^+/-^ mice showed no prefer in recognizing new object with a decreased discrimination index (Figure 1J and 1K), indicating ARIH1^+/-^ mice exhibited a decline in object recognition memory as well. To elucidate the specificity of ARIH1 deficiency on learning and memory, we also tested the changes of other behaviors in ARIH1^+/-^ mice such as locomotors activity, anxiety and depression, none of those tests showed significant difference in relation to WT mice (SI Figure 1D-1H).

**Figure 1.**
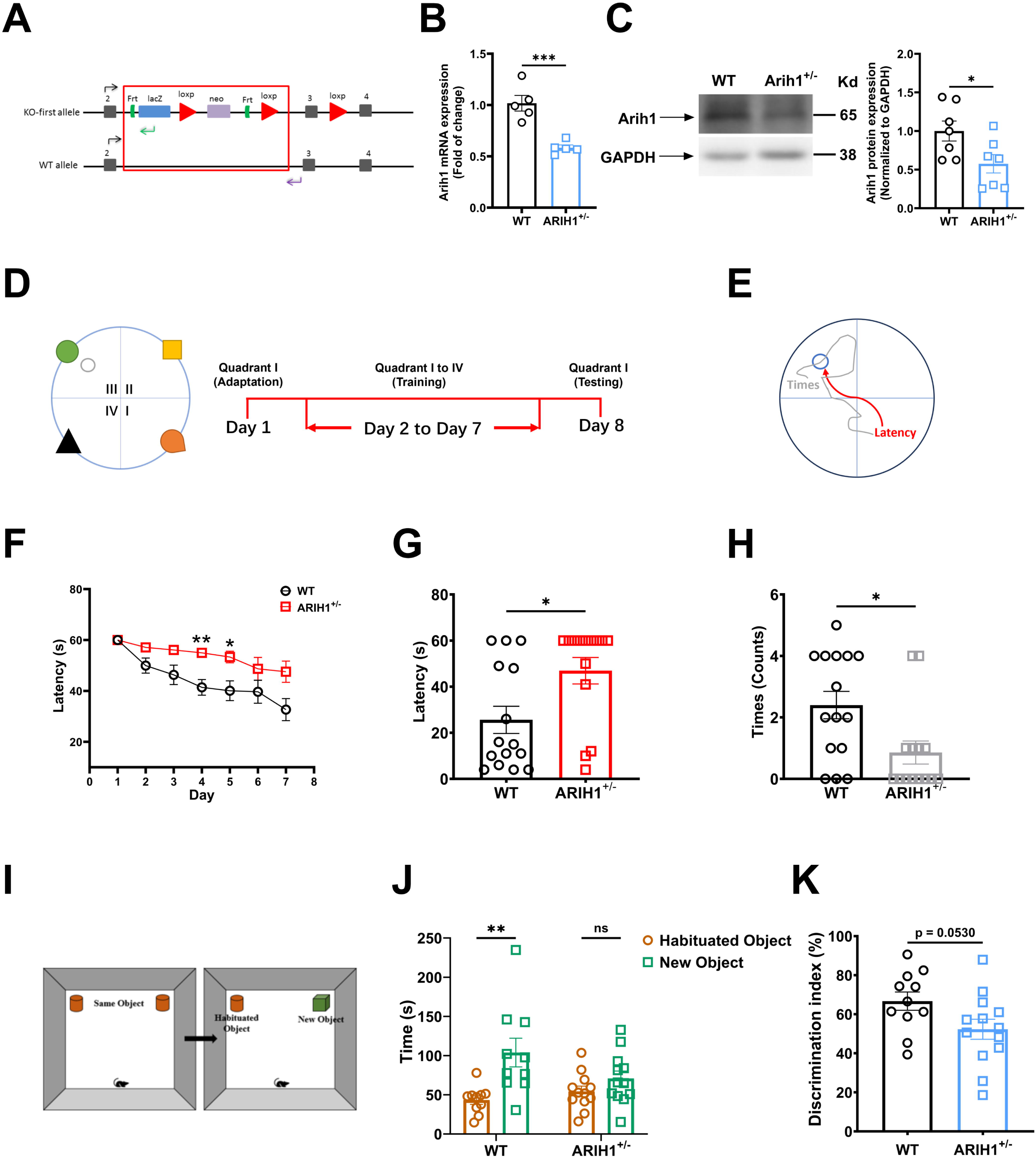
ARIH1^+/-^ mice exhibited declined learning and memory. (**A**) The embryonic stem cell vector information for constructing ARIH1 knockout mice. The hippocampal tissue of WT and ARIH1^+/-^ mice were dissected before the total RNA was prepared for qPCR (**B**) and whole lysis was prepared for immunoblot analysis (**C**) as described in the methods. The qPCR was probed by mouse ARIH1 primers, *n*=5. The immunoblot was probed by anti-ARIH1 antibody and showing blots are representative of at least 3 independent experiments. The mean intensity of bands was quantified using Image J and normalized to corresponding loading controls, *n*=7. (**D**, **E**) Schematic of Morris Water Maze (MWM) experiment. (**F**) The MWM experiment is composed of 3 sections which are adaptation (day 1), training (day 2 to 7), and testing (day 8). The latency on day 1 as well as the average latency (4 quadrants) on day 2 to 7 of male WT and ARIH1^+/-^ mice were calculated as described in the methods. During test on day 8, the latency (**G**) as well as times crossing the platform area (**H**) of male WT and ARIH1^+/-^ mice were recorded as described in the methods, *n*=14-15. (**I**) Schematic of Noval object recognition (NOR) experiment. (**J**, **K**) The NOR experiment is composed of training and testing sections, which are all videotaped for data analysis later. Both sections last 5 minutes and the interval between is 30 minutes, with 2 same objects placed in the arena during training, while 1 of the 2 objects was randomly replaced by another novel object during testing. Then, the object recognition time of male WT and ARIH1^+/-^ mice during testing was analyzed, and the discrimination index was calculated as described in the methods, *n*=11-13. * *p* < 0.05, *** *p* < 0.001, unpaired Student’s t test (**B**, **C**, **G**, **H**, **K**). * *p* < 0.05, ** *p* < 0.01, two-way ANOVA followed by Bonferroni’s test (**F**, **J**). Mean ± SEM.

### Local restoration the expression of ARIH1 in dorsal hippocampus is sufficient to rescue spatial memory impairment in ARIH1^+/-^ mice

Given dorsal hippocampus is a critical brain region in mediating spatial memory, and may be also partially involved in recognition memory, we thought to locally restore the expression of ARIH1 by injecting lentivirus expressing ARIH1 plasmids in ARIH1^+/-^ mice (Figure 2A and 2B) in dorsal hippocampus. As shown in Figure 2C-2E, dorsal hippocampal ARIH1 complement significantly rescued the spatial memory decline in ARIH1^+/-^ mice. In addition, dorsal hippocampal ARIH1 complement showed a slight recovery on the novel object recognition and discrimination index in ARIH1^+/-^ mice although the increase didn’t reach a statistic difference (Figure 2F and 2G). This data further confirmed the importance of ARIH1 in learning and memory.

**Figure 2.**
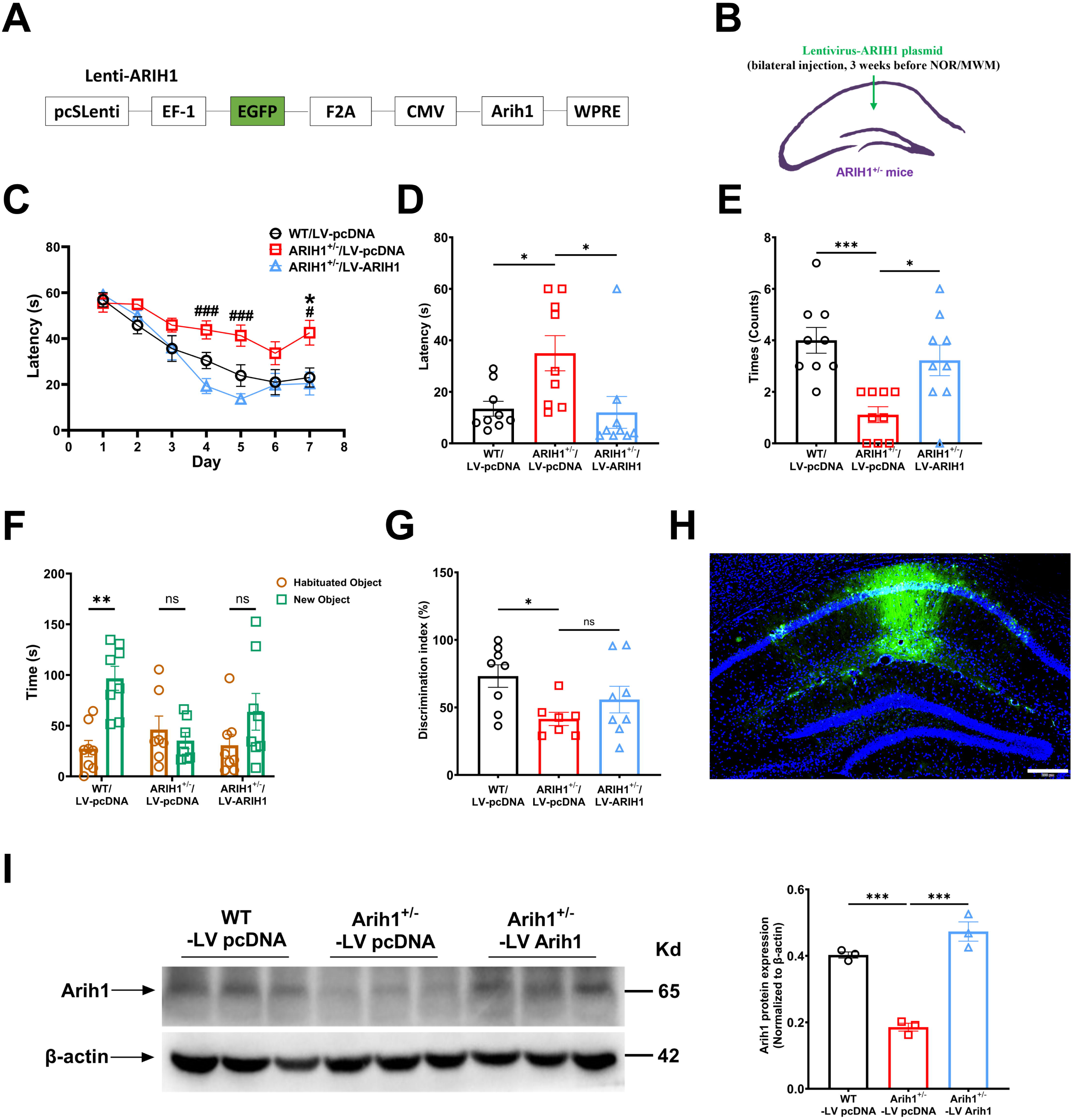
Dorsal hippocampal expression of ARIH1 restored the spatial memory decline in ARIH1^+/-^ mice. Schematic of the lentivirus expressing ARIH1 plasmid (**A**) and brain injection site (**B**) in mice. (**C**) The MWM experiment is composed of 3 sections which are adaptation (day 1), training (day 2 to 7), and testing (day 8). The latency on day 1 as well as the average latency (4 quadrants) on day 2 to 7 of WT and ARIH1^+/-^ mice, injected with lenti-vector or lenti-ARIH1 plasmid, were calculated as described in the methods. *, ARIH1^+/-^/LV-pcDNA vs. WT/LV-pcDNA; # and ###, ARIH1^+/-^/LV-pcDNA vs. ARIH1^+/-^/LV-ARIH1. During test on day 8, the latency (**D**) as well as times crossing the platform area (**E**) of WT and ARIH1^+/-^ mice, injected with lenti-vector or lenti-ARIH1 plasmid, were recorded as described in the methods, *n*=9. (**F**, **G**) The NOR experiment is composed of training and testing sections, which are all videotaped for data analysis later. Both sections last 5 minutes and the interval between is 30 minutes, with 2 same objects placed in the arena during training, while 1 of the 2 objects was randomly replaced by another novel object during testing. Then, the object recognition time of WT and ARIH1^+/-^ mice, injected with lenti-vector or lenti-ARIH1 plasmid during testing was analyzed, and the discrimination index was calculated as described in the methods, *n*=7-8. (**H**) Representative image for showing the site of injection. Scale bar, 1 mm. (**I**) The dorsal hippocampal tissue of WT and ARIH1^+/-^ mice, injected with lenti-vector or lenti-ARIH1 plasmid, were dissected and whole lysis was prepared for immunoblot analysis probed by anti-ARIH1 antibody. The mean intensity of bands was quantified using Image J and normalized to corresponding loading controls, *n*=3. * *p* < 0.05, *** *p* < 0.001, *ns*. not significant, one-way ANOVA followed by Bonferroni’s test (**D**, **E**, **G**) or by Tukey’s test (**I**). * *p* < 0.05, ** *p* < 0.01, ### *p* < 0.001, *ns*. not significant, two-way ANOVA followed by Bonferroni’s test (**C**, **F**). Mean ± SEM.

After completing the behavioral examinations, the brain tissue was subjected to immunohistochemistry and immunoblot analysis to verify injection site of the lentivirus and the expression of ARIH1 in the dorsal hippocampus. The results are as expected and shown in (Figure 2H and 2I).

### Elevated dorsal hippocampal GIRK2 expression may contribute to the spatial learning and memory decline in ARIH1^+/-^ mice

In order to explore the mechanism for ARIH1-regulated learning and memory, the hippocampal tissues of ARIH1^+/-^ mice and WT control mice were subjected to label-free quantitative proteomics analysis (Figure 3A). Out of 4448 proteins identified, the expression of 139 proteins in ARIH1^+/-^ mice were significantly increased as compared to that of WT controls, whilst other 66 proteins were deceased (Figure 3B and 3C).

**Figure 3.**
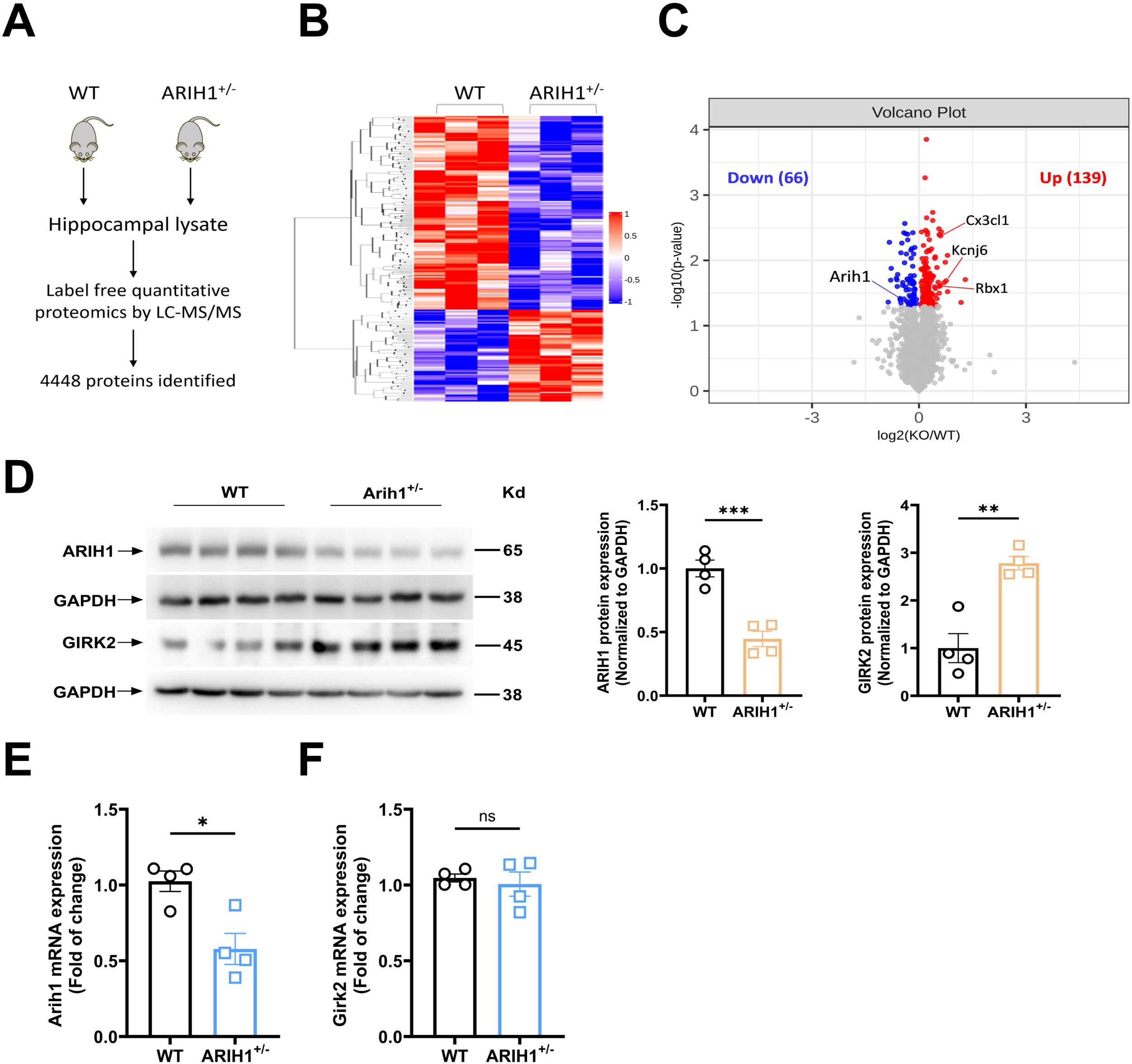
GIRK2 expression was elevated in the dorsal hippocampus of ARIH1^+/-^ mice. (**A**) Workflow of quantitative proteomic analysis. Proteomic data were obtained from mice hippocampus, *n*=3. (**B**) Heatmaps of differentially expressed (DE) proteins in WT and ARIH1^+/-^ hippocampal samples. (**C**) Volcano plot for DE proteins (139 upregulated, 66 downregulated) in ARIH1^+/-^ hippocampal samples compared with WT hippocampus. Red and blue dots indicate statistical significance DE proteins. The hippocampal tissue of WT and ARIH1^+/-^ mice were dissected and whole lysis was prepared for immunoblot analysis (**D**) and total RNA was prepared for qPCR analysis (**E**, **F**) as described in the methods. The qPCR was probed by mouse ARIH1 or GIRK2 primers, *n*=4. The immunoblot was probed by anti-ARIH1 or anti-GIRK2 antibody. The mean intensity of bands was quantified using Image J and normalized to corresponding loading controls, *n*=4. * *p* < 0.05, ** *p* < 0.01, *** *p* < 0.001, *ns*. not significant, unpaired Student’s t test (**D**, **E**, **F**). Mean ± SEM.

Among the increased proteins, 10 proteins are more than 1.5fold higher including Kcnj6 (Figure 3C). Kcnj6 gene encodes G protein-gated inwardly rectifying K^+^ channels subunit 2 (GIRK2). GIRK2 is highly expressed in rodent brain which is known to regulate learning and memory. We deduced that GIRK2 might be a substrate of ARIH1. We thus first confirmed that the elevated protein expression of GIRK2 is not attributed to the transcriptional increase in mRNA expression of GIRK2 since qPCR did not detect a significant change of GIRK2 in ARIH1^+/-^ mice as compared to that of controls (Figure 3D-3F). Suggesting that the elevated GIRK2 protein is likely attributed to attenuation in protein turnover, but not the transcription.

To explore the functional role of GIRK2 protein upregulation in ARIH1^+/-^ mice, we employed the widely used GIRK current inhibitor Tipepidine [28] to test its effect on the impaired learning and memory in ARIH1^+/-^ mice. As shown in Figure 4C-4E, Tipepidine administration (*i.p.*, 20mg/kg) successfully rescued the spatial learning and memory decline in ARIH1^+/-^ mice, the treatment did not significantly alter the basal spatial learning and memory in WT mice (SI Figure 2A-2C). In the contrast, Tipepidine administration didn’t improve the recognition memory deficit of ARIH1^+/-^ mice as evidenced in NOR test (Figure 4F and 4G), which may imply that there are other mechanisms underlying the recognition memory decline of ARIH1^+/-^ mice. In support, local administration of a specific GIRK channel inhibitor Tertiapin-Q (*i.c.*, 0.5μl, 0.25mM) [29, 30] also produced the similar results (Figure 4H-4J). In agreement with observations in Tipepidine administration, Tertiapin-Q failed to rescue the declined recognition memory of ARIH1^+/-^ mice as well (Figure 4K and 4L).

**Figure 4.**
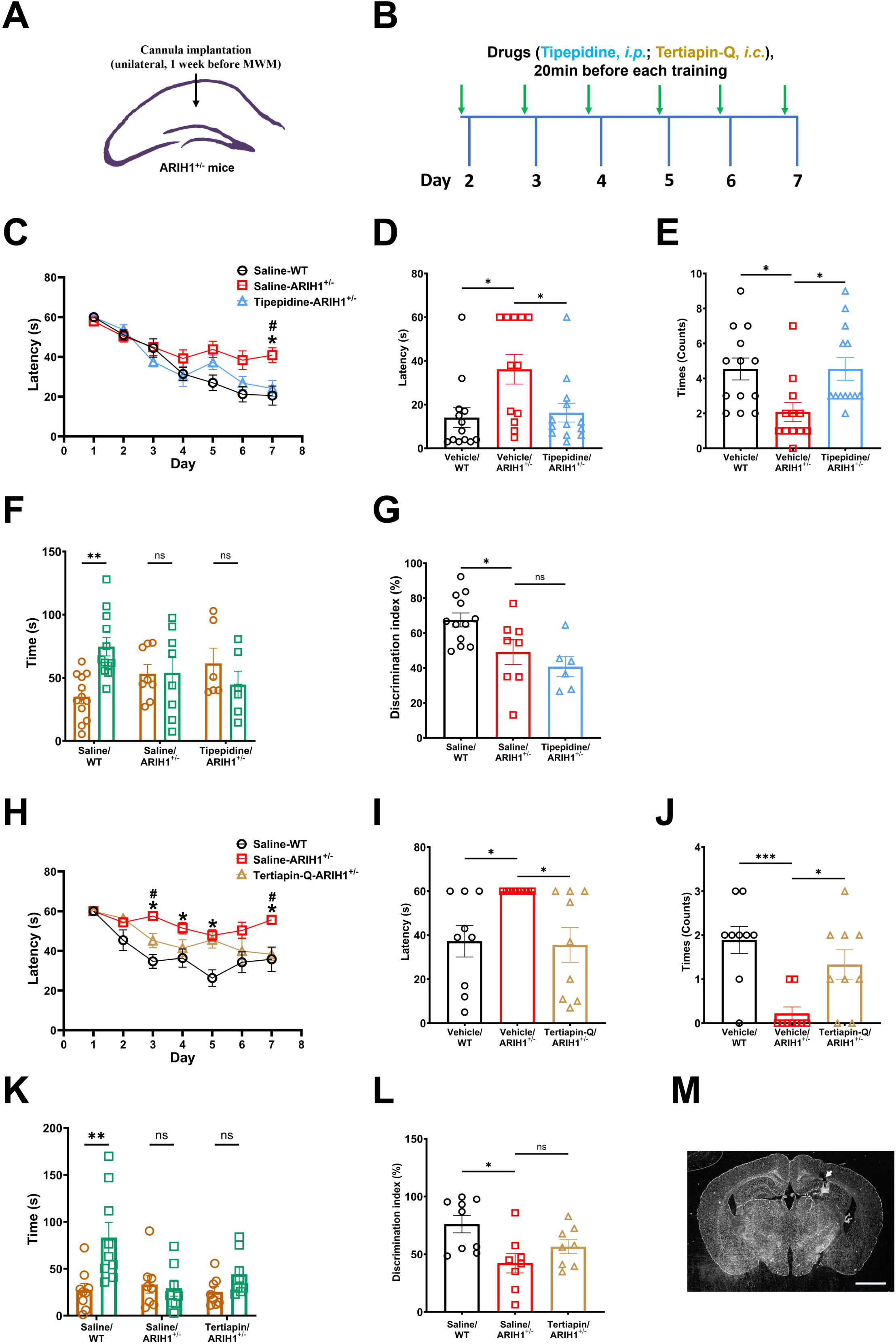
Elevated dorsal hippocampal GIRK2 expression may contribute to the spatial learning and memory decline in ARIH1^+/-^ mice. (**A**) Schematic of cannular implantation site. (**B**) Timeline for drug administration. (**C**) The MWM experiment is composed of 3 sections which are adaptation (day 1), training (day 2 to 7), and testing (day 8). The latency on day 1 as well as the average latency (4 quadrants) on day 2 to 7 of WT and ARIH1^+/-^ mice, administrated with Saline or Tipepidine (*i.p.*, 20mg/kg), were calculated as described in the methods. *, Saline-ARIH1^+/-^ vs. Saline-WT; #, Saline-ARIH1^+/-^ vs. Tipepidine-ARIH1^+/-^. During test on day 8, the latency (**D**) as well as times crossing the platform area (**E**) of WT and ARIH1^+/-^ mice, administrated with Saline or Tipepidine, were recorded as described in the methods, *n*=12-13. (**F**, **G**) The NOR experiment is composed of training and testing sections, which are all videotaped for data analysis later. Both sections last 5 minutes and the interval between is 30 minutes, with 2 same objects placed in the arena during training, while 1 of the 2 objects was randomly replaced by another novel object during testing. Then, the object recognition time of WT and ARIH1^+/-^ mice, administrated with Saline or Tipepidine (*i.p.*, 20mg/kg) during testing was analyzed, and the discrimination index was calculated as described in the methods, *n*=6-12. (**H**) The latency on day 1 as well as the average latency (4 quadrants) on day 2 to 7 of WT and ARIH1^+/-^ mice, administrated with Saline or Tertiapin-Q (*i.c.*, 0.5μl, 0.25mM), were calculated as described in the methods. *, Saline-ARIH1^+/-^ vs. Saline-WT; #, Saline-ARIH1^+/-^ vs. Tertiapin-Q-ARIH1^+/-^. During test on day 8, the latency (**I**) as well as times crossing the platform area (**J**) of WT and ARIH1^+/-^ mice, administrated with Saline or Tertiapin-Q, were recorded as described in the methods, *n*=9. (**K**, **L**) The object recognition time of WT and ARIH1^+/-^ mice, administrated with Saline or Tertiapin-Q (*i.c.*, 0.5μl, 0.25mM) during testing was analyzed, and the discrimination index was calculated as described in the methods, *n*=8-9. (**M**) Representative image for showing the site of cannular implantation. Scale bar, 1 mm. * *p* < 0.05, *** *p* < 0.001, *ns*. not significant, one-way ANOVA followed by Bonferroni’s test (**D**, **E**, **G**, **I**, **J**, **L**). * *p* < 0.05, ** *p* < 0.01, # *p* < 0.05, *ns*. not significant, two-way ANOVA followed by Bonferroni’s test (**C**, **F**, **H**, **K**). Mean ± SEM.

Taken together, our data revealed that increased GIRK2 expression in dorsal hippocampus is likely responsible for the declined spatial learning and memory of ARIH1^+/-^ mice.

### ARIH1 deficiency damages the ubiquitination and degradation of GIRK2

We next asked how ARIH1 defect altered GIRK2 turnover. We constructed ARIH1 knockdown HT-22 cells through transduction lentivirus expressing specific ARIH1 shRNA. As expected, knockdown of ARIH1 by shRNA (948-HT-22, 949-HT-22 cells) increased the expression of GIRK2 proteins (Figure 5A, 5A_1_ and 5A_2_), which replicated the observation in hippocampus in ARIH1^+/-^ mice. Moreover, re-expression of ARIH1 in 948-HT-22 or 949-HT-22 cells by transfection ARIH1 plasmids significantly decreased GIRK2 protein expression to the control level (Figure 5B and 5B_1_-5B_4_). In consistent with data from brain tissues of ARIH1^+/-^ mice, knockdown of ARIH1 in both 948-HT-22 and 949-HT-22 cells did not alter mRNA expression of GIRK2 (Figure 5D and 5E), further demonstrated that altered protein degradation may be involved in GIRK2 upregulation in response to ARIH1 deficiency. Indeed, as shown in Figure 5C and 5C_1_, the ubiquitination of GIRK2 in ARIH1 knockdown cells was dramatically suppressed, confirming that ARIH1 defect-induced GIRK2 upregulation was indeed attributed to the damaged ubiquitination.

**Figure 5.**
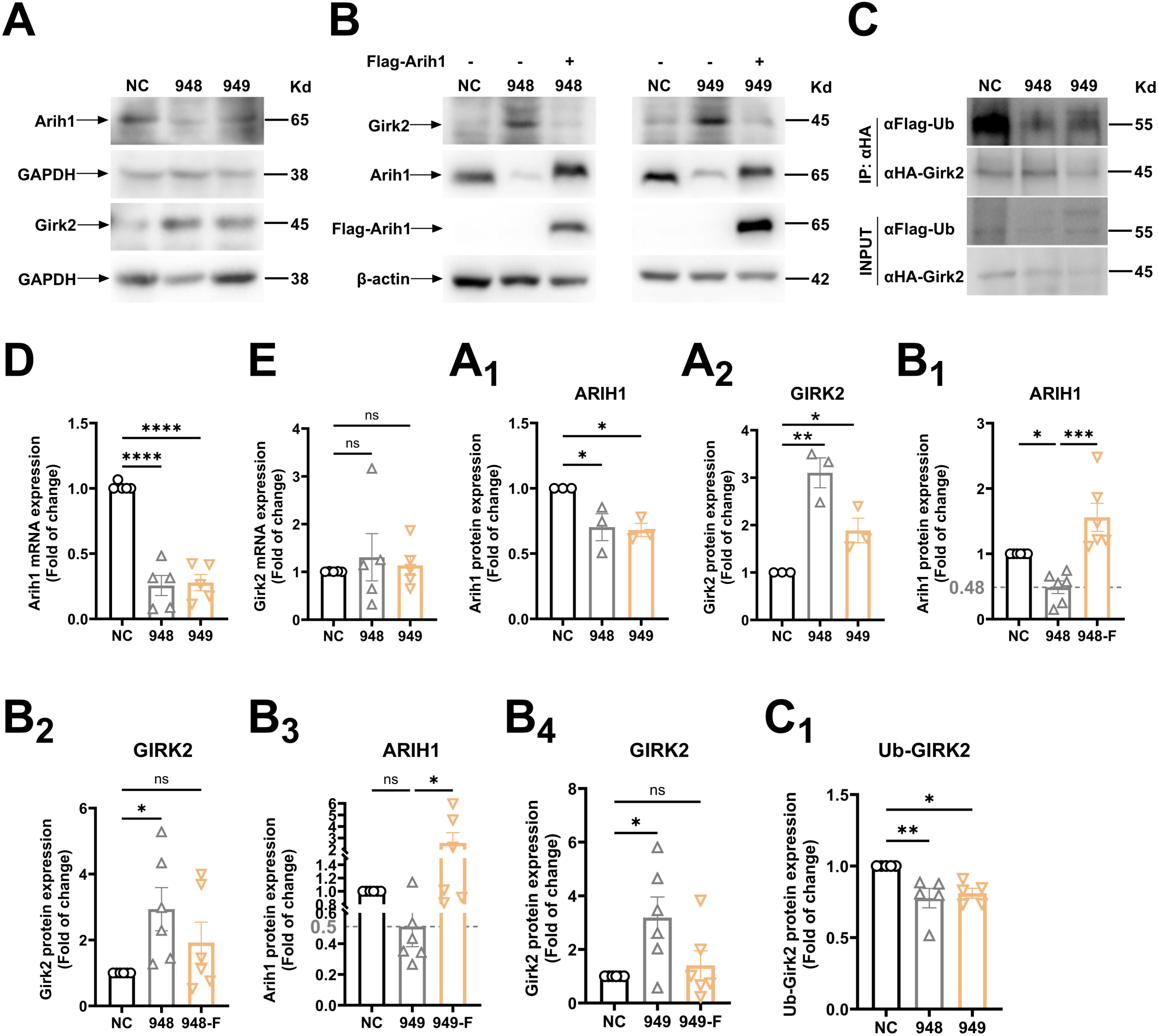
ARIH1 mediated the ubiquitination and degradation of GIRK2. HT-22 cells were infected with lentivirus expressing scramble (NC) or Arih1-shRNA of two different on-target sequence (948, 949). NC and Arih1 knockdown cells were selected and maintained in cell culture media supplemented with puromycin (2.5μg/ml for selection, and 1μg/ml for maintaining) as described in the methods. (**A**) Whole cell lysates were prepared from HT-22 NC and Arih1 knockdown cells for immunoblot analysis probed by anti-Arih1 or anti-Girk2 antibodies. (**B**) Arih1 knockdown cells were transfected with vector or Flag-Arih1 plasmids. After 48hours transfection, whole cell lysates were prepared for immunoblot analysis probed by anti-Girk2, anti-Arih1, or anti-Flag antibodies. (**C**) HT-22 NC and Arih1 knockdown cells were transfected with Flag-Ub and HA-Girk2 plasmids. After 48hours transfection, whole cell lysates were prepared and immunoprecipitation was performed to examine the ubiquitination of Girk2 using anti-Flag or anti-HA antibodies. Showing blots are representative of at least 3 independent experiments. The mean intensity of bands was quantified using Image J and normalized to corresponding loading controls (**A_1_-A_2_**, **B_1_-B_4_**), or to corresponding input loadings (**C_1_**). (**D**, **E**) The total RNA was prepared from HT-22 NC and Arih1 knockdown cells for qPCR analysis with mouse ARIH1 or GIRK2 primers, *n*=5. * *p* < 0.05, ** *p* < 0.01, *** *p* < 0.001, **** *p* < 0.0001, *ns*. not significant, one-way ANOVA followed by Tukey’s test, *n*=3-6. Mean ± SEM.

### Selective knockdown ARIH1 in dorsal hippocampal CaMKII-expressing neurons mimics the impairment of learning and memory in ARIH1^+/-^ mice

To elucidate the specific neurons that mediate ARIH1 deficient-induced impairment of learning and memory in mice. We employed CaMKII-Cre^+^ mice by local injecting AAV-DIO-ARIH1 shRNA (sequence of 948) into the dorsal hippocampus to selectively knockdown ARIH1 in the neurons. Three weeks after AAV injection, the MWM test was then conducted (Figure 6A and 6B). As shown in Figure 6C-6E, specific knockdown ARIH1 in mice dorsal hippocampal CaMKII-positive neurons markedly damaged their spatial learning and memory in comparison with mice subjected to vector injections. The efficiency and specificity of knockdown was confirmed by fluorescence image and immunoblot (Figure 6F and 6G). Immunofluorescent analysis verified the co-expression of AAV-DIO-ARIH1 shRNA with CaMKII positive neurons, but not with GFAP or IB1 positive cells (Figure 6H). As expected, ARIH1 knockdown in dorsal hippocampal CaMKII-expressing neurons robustly increased local expression of GIRK2 (Figure 6I and 6J).

**Figure 6.**
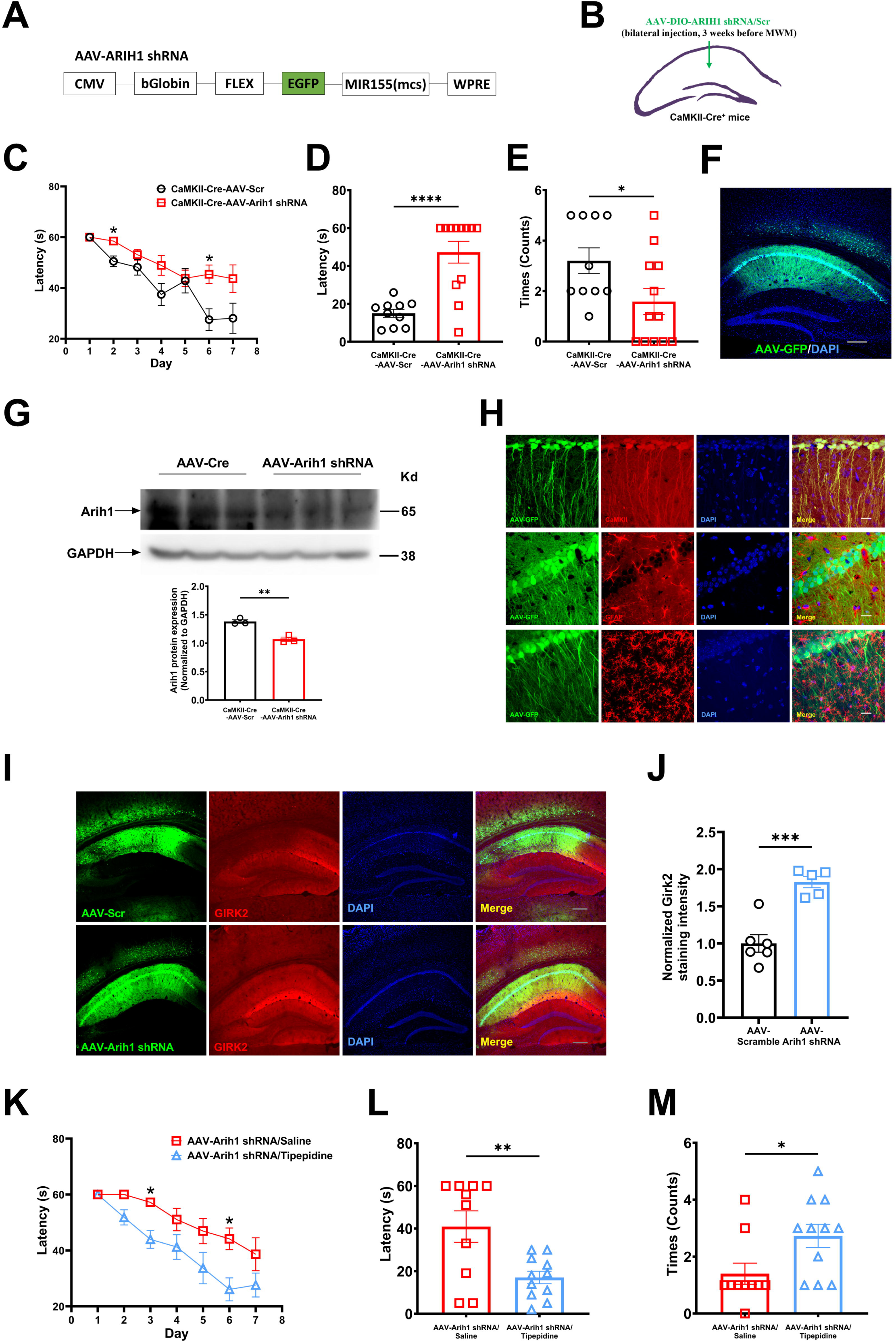
Elevated GIRK2 expression may contribute to the spatial memory decline of mice with ARIH1 knockdown in dorsal hippocampal CaMKII-expressing neurons. Schematic of the AAV-ARIH1 shRNA (**A**) and brain injection site (**B**) in mice. (**C**) The MWM experiment is composed of 3 sections which are adaptation (day 1), training (day 2 to 7), and testing (day 8). The latency on day 1 as well as the average latency (4 quadrants) on day 2 to 7 of AAV-Scramble or AAV-ARIH1 shRNA injected mice were calculated as described in the methods. During test on day 8, the latency (**D**) as well as times crossing the platform area (**E**) of AAV-Scramble or AAV-ARIH1 shRNA injected mice were recorded as described in the methods, *n*=10-12. (**F**) Representative image for showing the site of injection. Scale bar, 100μm. (**G**) The dorsal hippocampal tissue of AAV-Scr and AAV-ARIH1 shRNA injected mice were dissected and whole cell lysates were prepared for immunoblot analysis probed by anti-ARIH1 antibody. The mean intensity of bands was quantified using Image J and normalized to corresponding loading controls, *n*=3. (**H**) Representative image for showing co-expression of AAV-GFP with CaMKII, but not with GFAP or IB1. Scale bar, 20μm. (**I**) Representative image of immunostaining of Girk2 in mice with dorsal hippocampal injection of AAV-Scramble or AAV-ARIH1 shRNA. Scale bar, 100μm. (**J**) Quantification of the intensity of Girk2 staining, *n*=5. (**K**) The latency on day 1 as well as the average latency (4 quadrants) on day 2 to 7 of AAV-ARIH1 shRNA injected mice, administrated with Saline or Tipepidine (*i.p.*, 20mg/kg), were calculated as described in the methods. During test on day 8, the latency (**L**) as well as times crossing the platform area (**M**) of AAV-ARIH1 shRNA injected mice, administrated with Saline or Tipepidine, were recorded as described in the methods, *n*=10-11. * *p* < 0.05, ** *p* < 0.01, *** *p* < 0.001, **** *p* < 0.0001, unpaired Student’s t test (**D**, **E**, **G**, **J**, **L**, **M**). * *p* < 0.05, two-way ANOVA followed by Bonferroni’s test (**C**, **K**). Mean ± SEM.

In contrast, we found that in transgenic mice with ARIH1 conditional knockdown in either Parvalbumin^+^ (PV, PV-ARIH1-KO) or Somatostatin^+^ (SST, SST-ARIH1-KO) neurons exhibited no difference in MWM test in comparison with WT controls (SI Figure 3A-3C, 3F-3H). The knockdown efficiency of ARIH1 in PV- or SST-expressing neurons was confirmed by immunofluorescent analysis (SI Figure 3D-3E and 3I-3J). These observations indicated that CaMKII-expressing neurons is the major subtype of neurons to mediate ARIH1 deficiency-impaired learning and memory in ARIH1^+/-^ mice.

To verify the functional role of GIRK2 upregulation of CaMKII-expressing neurons in spatial learning and memory, the GIRK inhibitor tipepidine was administrated in mice with selective ARIH1 knockdown in dorsal hippocampal CaMKII-expressing neurons. As shown in Figure 6K-6M, tipepidine administration significantly improved the spatial learning and memory.

Taken together, our data provided the first evidence that ARIH1 in dorsal hippocampal CaMKII-expressing neurons is critical in mediating spatial learning and memory via modulating the expression of GIRK2.

## Discussion

Recent studies have revealed roles of ARIH1 in chemotherapy resistance [3], cancer progression [4], as well as immunoregulation [5–8] in rodents. In humans, variants in ARIH1 are found to be associated with developing aortic aneurysms [9]. In the current study, we found that ARIH1-mutant mice (ARIH1^+/-^) exhibited a declined spatial and recognition learning and memory. We provided the first evidence that ARIH1 deficiency induced the enhanced expression of GIRK2 in ARIH1^+/-^ mice. Moreover, local knockdown of ARIH1 in dorsal hippocampal CaMKII-expressing neurons was sufficient to mimic the impairment of spatial learning and memory, which was not the case if knockdown of ARIH1 in PV^+^ or SST^+^ neurons. These observations indicated that CaMKII-expressing hippocampal neurons played essential roles in ARH1 deficiency-induced damage in spatial learning and memory. Furthermore, application of GIRK2 inhibitor effectively restored the impaired spatial learning and memory of mice with ARIH1 knockdown of CaMKII-expressing neurons. Lastly, we demonstrated *in vitro* that knockdown of ARH1 in HT 22 neuronal cells resulted in the elevated GIRK2 expression which was not attributed to the enhanced synthesis but rather due to the impaired ubiquitination that ultimately led to the suppression of GIRK2 degradation. The present data thus provided the first evidence to uncover the important role of ARIH1 in mediating learning and memory via upregulating GIRK2 expression in mice dorsal hippocampal CaMKII-expressing neurons.

ARIH1 gene is the human homolog of *Drosophila* ariadne-1 (Ari-1). Survivors of Ari-1 null alleles exhibited motor impairments [11], however, ARIH1^+/-^ mice exhibited normal basal locomotor activity and basal performance in rotarod and pole tests as compared to control mice. One possible explanation for these discrepancies could be the differences in ARIH1 protein levels since the decrease of ARIH1 in Ari-1 null flies is more robust [11]. Interestingly, we observed a dramatic impaired learning and memory in ARIH1^+/-^ mice in both MWM and NOR tests. This is the first evidence to reveal the role of ARIH1 in learning and memory, although previous studies have shown the modulatory effect of a number of other E3 ligases in learning and memory [17, 18]. Recent study demonstrated that ARIH1 works in unison with Cullin-RING ubiquitination ligases (CRLs), especially CRL1-3 and CRL4A, to regulate substrate ubiquitination [31]. CRL1 and CRL2 are involved in neuronal pruning and the development of ALS (Amyotrophic Lateral Sclerosis) [32, 33], respectively, while knockdown the autism risk gene Cul3 induced hyperlocomotion, anxiety, social and cognitive defect, as well as decreased morphine dependence [34–38]. Thus, future studies would be interesting to examine the role and underlying mechanisms of ARIH1 in these CRLs related neurological phenotypes.

The consent that dorsal hippocampus is an essential brain region for spatial learning and memory is well established, whilst the role of dorsal hippocampal dependency for recognition memory is controversy. For example, transiently inactivate the dorsal hippocampus of mice by local microinjection of muscimol (GABA_A_ agonist) or by the chemogenetic approach of designer receptor exclusively activated by designer drugs (DREADDs), robustly impaired the recognition memory [39, 40]. In contrast, the recognition memory in rats with permanent lesion of the hippocampus by local injection of NMDA (excitotoxin) remains intact [41, 42]. These discrepancies may arise due to post-lesion compensatory plasticity within neural circuits since rats with bilateral excitotoxic lesions of the hippocampus is normal in anterograde object recognition test [43]. Nevertheless, lesion of perirhinal cortex (PC) strongly declined the recognition memory of rodent and non-human primates [41, 44], while medial prefrontal cortex (mPFC) in rodents is required for recognition of an object in a particular context [45], suggesting the involvement of hippocampus independent mechanisms for recognition memory. We found that local restoration the expression of ARIH1 in dorsal hippocampus is sufficient to rescue the spatial learning and memory impairment in ARIH1^+/-^ mice, which is in agreement with the notion that dorsal hippocampus is essential for spatial learning and memory. In contrast, dorsal hippocampal ARIH1 complement only slightly recovered the novel object recognition and discrimination index in ARIH1^+/-^ mice, suggesting an involvement of other brain regions, such as PC or mPFC, in mediating the recognition memory decline of ARIH1^+/-^ mice. Further studies are needed to examine this hypothesis.

GIRK family is composed of 4 subunits GIRK1-4. GIRK2 is highly expressed in neocortex, cerebellum, and hippocampus existing as homo-tetramer and hetero-tetrameric complexes with other subunits in modulating neuronal excitability [46]. Several lines of evidence reveal that an optimal range of GIRK2 activity is critical for normal synaptic plasticity. For instance, GIRK2 null mutation mice or wild type mice applied with GIRK channel blocker [47], as well as Kcnj6 gene (encoding GIRK2) triploid mice that with elevated GIRK2 expression, exhibit hampered depotentiation of LTP and accelerated LTD in their hippocampal neurons [48]. Importantly, both loss- and gain-of-function studies for GIRK2 channel indicate a close relationship between dysregulation of GIRK2 channel and cognitive defect. For example, pharmacological activation of GIRK2 rescued memory decline in Alzheimer’s Disease model mice, where the transcription of GIRK2 decreased [49–51], while Kcnj6 gene triploid mice exhibited declined hippocampal-dependent learning and memory, and the increased GIRK2 transcription may largely contribute to the cognitive impairment in Down syndrome model mice [48, 52]. Moreover, patients with either gain-of-function mutation or loss-of-function mutation of Kcnj6, which causes the Keppen-Lubinsky Syndrome exhibit severe developmental delay and intellectual disability [53, 54]. Through proteomic analysis and immunoblot verification, we observed an upregulation of GIRK2 protein in the hippocampus of ARIH1^+/-^ mice comparing with the WT controls. Interestingly, administration of GIRK inhibitors Tipepidine systematically or Tertiapin-Q in the dorsal hippocampus successfully ameliorated the spatial learning and memory decline of ARIH1^+/-^ mice. Our data is in agreement with a recent report in which knockout the GIRK2 in forebrain could impair hippocampal-dependent memory in mice [55]. Indeed, the apparent inconsistency between our observations and previous reports could be explained by another investigation showed that either overactive or inhibiting dorsal hippocampal GIRK2 could result in a declined learning and memory in mice [30], implying that GIRK2 homeostasis is critical important. It is worthy noticing that intracerebroventricular injection of either GIRK2 activator ML297 or inhibitor Tertiapin-Q impaired the recognition memory of the mice in NOR test, which indicates an involvement of GIRK channel-dependent mechanism for the recognition memory [30]. In this study, we revealed that systematical administration of Tipepidine successfully rescued declined spatial learning and memory, but not recognition memory of ARIH1^+/-^ mice, suggesting that ARIH1 deficiency may impair the recognition memory through other mechanisms rather than the upregulation of GIRK2.

Another important finding of the present study is that we revealed GIRK2 could be the substrate of ARIH1 for ubiquitination and degradation. Function as an E3 ligase, ARIH1 mediates both non-proteolytic ISG15 modification (Interferon-stimulated gene 15, ISGylation) and proteasome-dependent proteolytic ubiquitination modification on its substrates. For example, ARIH1 catalyzes the mono-ISGylation and induces the oligomerization of cyclic GMP-AMP synthase (cGAS), thereby promoting antiviral immunity and autoimmunity [6], while ARIH1 also promotes anti-tumor immunity by targeting PD-L1 for proteasomal degradation [5]. We found that knockdown ARIH1 robustly upregulated the protein expression of GIRK2 *in vivo* and *in vitro* without alteration of the protein synthesis, while complement expressing ARIH1 in ARIH1-knockdown cells normalized GIRK2 protein to the control level. Moreover, we observed an inhibition of GIRK2 ubiquitination in ARIH1-knockdown cells. Since the ARIH1 works in unison with CRLs to regulate substrate ubiquitination [31], further studies are warranted to test any potential involvement of CRLs in regulating GIRK2 ubiquitination and degradation.

Moreover, we also identified that the dorsal hippocampal CaMKII-expressing neurons are critical in mediating the declined spatial learning and memory in ARIH1-deficient mice. The hippocampal CaMKII-expressing pyramidal neurons are well-documented pivotal players for spatial cognitive functions. The number of mushroom spines and synapses on pyramidal neurons increases with learning and memory in the MWM [56], while inactivation or genetic knockout NMDA receptors of dorsal hippocampal pyramidal neurons, which impairs local LTP, robustly declined spatial learning and memory in mice [57, 58]. We observed that selective knockdown dorsal hippocampal ARIH1 of CaMKII-expressing neurons by AAV injection mimics the impairment of spatial learning and memory in ARIH1^+/-^ mice. In addition to pyramidal neurons, gamma-aminobutyric acid (GABAergic)-expressing, especially PV^+^ and SST^+^ interneurons, are also important in hippocampal-dependent learning and memory. The PV^+^ neurons mediate learning and memory in a task specific manner, as mice lacking NMDA receptors or neuroligin-3 in PV^+^ neurons exhibited disrupted gamma oscillations and impaired fear conditioning and memory, as well as declined spatial working and memory in Y-maze and T-maze tests, but maintaining intact spatial learning and memory in MWM [59–61]. In consistent, we observed that PV-ARIH1-KO mice exhibited normal spatial learning and memory in MWM. Moreover, direct inactivating SST^+^ neurons or blunting synaptic plasticity in SST^+^ neurons through conditional down regulating mTORC1 (mammalian target of rapamycin complex 1) of SST^+^ neurons resulted in the impaired fear conditioning and memory, as well as declined spatial memory in Barnes maze, whilst their spatial learning remains intact [62, 63]. In the present study, we observed that SST-ARIH1-KO mice exhibited normal spatial learning and memory in MWM. The discrepancy between our data and previous report in terms of spatial memory may arise due to the differences in the tasks that has been used, as serotonin transporter knockout mice or 3,4-methylenedioxymethamphetamine-treated mice exhibited altered spatial task performance in MWM but not in Barnes maze [64, 65]. Alternatively, future studies are needed to examine the synaptic plasticity integrity of SST^+^ neurons in SST-ARIH1-KO mice, which may help to interpret the discrepancies.

In summary, our findings uncovered the role of ARIH1 in learning and memory. Moreover, we demonstrated that ARIH1 deficiency resulted in upregulation of GIRK2 due to impaired ubiquitination and degraded. Furthermore, we elucidated that dorsal hippocampal CaMKII-expressing neurons is main subtype of neurons that mediates the spatial learning and memory decline in ARIH1-deficient mice.

## Acknowledgments

We acknowledge CAM-SU Genomic Resource Center, Soochow University, National Center for International Research of Genomics (2017B01012) and 2018YFA0801100 “Establishment of mouse developmental and metabolic phenotype repository”.

**SI Figure 1. ARIH1^+/-^ female mice exhibited declined spatial memory, and ARIH1^+/-^ male mice behaved normal in locomotion and anxiety**. (**A**) The MWM experiment is composed of 3 sections which are adaptation (day 1), training (day 2 to 7), and testing (day 8). The latency on day 1 as well as the average latency (4 quadrants) on day 2 to 7 of female WT and ARIH1^+/-^ mice were calculated as described in the methods. During test on day 8, the latency (**B**) as well as times crossing the platform area (**C**) of female WT and ARIH1^+/-^ mice were recorded as described in the methods, *n*=11-12. For male WT and ARIH1^+/-^ mice, the locomotor activity was recorded for 30 minutes (**D**), the motor function was examined in rotarod (**E**) and pole test (**F**), the anxiety behavior was examined in elevated plus maze test (**G**), and the depressive behavior was examined in tail suspension test as described in the methods (**H**), *n*=10-14. * *p* < 0.05, ** *p* < 0.01, *ns*. not significant, unpaired Student’s t test (**B-H**). ** *p* < 0.01, *** *p* < 0.001, two-way ANOVA followed by Bonferroni’s test (**A**). Mean ± SEM.

**SI Figure 2. Effect of Tipepidine on MWM of WT mice**. (**A**) The MWM experiment is composed of 3 sections which are adaptation (day 1), training (day 2 to 7), and testing (day 8). The latency on day 1 as well as the average latency (4 quadrants) on day 2 to 7 of WT mice, administrated with Saline or Tipepidine (*i.p.*, 20mg/kg), were calculated as described in the methods. During test on day 8, the latency (**B**) as well as times crossing the platform area (**C**) of WT mice, administrated with Saline or Tipepidine, were recorded as described in the methods, *n*=8-9. *ns*. not significant, unpaired Student’s t test (**B**, **C**). Mean ± SEM.

**SI Figure 3. ARIH1 Parvalbumin (PV)- or Somatostatin (SST)-expressing neuron conditional knockout mice exhibited normal spatial memory**. (**A**) The MWM experiment is composed of 3 sections which are adaptation (day 1), training (day 2 to 7), and testing (day 8). The latency on day 1 as well as the average latency (4 quadrants) on day 2 to 7 of ARIH1^f/f^ and PV-ARIH1^f/f^ mice were calculated as described in the methods. During test on day 8, the latency (**B**) as well as times crossing the platform area (**C**) of ARIH1^f/f^ and PV-ARIH1^f/f^ mice were recorded as described in the methods, *n*=5-6. (**D**) Representative image of PV and ARIH1 double-immunostaining of ARIH1^f/f^ and PV-ARIH1^f/f^ mice. Scale bar, 50μm. (**E**) Fiji software was used to quantify the immunofluorescence intensity of ARIH1 staining co-expressed with PV staining positive cells, *n*= 10-11. (**F**) The latency on day 1 as well as the average latency (4 quadrants) on day 2 to 7 of ARIH1^f/f^ and SST-ARIH1^f/f^ mice were calculated as described in the methods. During test on day 8, the latency (**G**) as well as times crossing the platform area (**H**) of ARIH1^f/f^ and SST-ARIH1^f/f^ mice were recorded as described in the methods, *n*=9-10. (**I**) Representative image of SST and ARIH1 double-immunostaining of ARIH1^f/f^ and SST-ARIH1^f/f^ mice. Scale bar, 50μm. (**J**) Fiji software was used to quantify the immunofluorescence intensity of ARIH1 staining co-expressed with SST staining positive cells, *n*= 10-11. * *p* < 0.05, **** *p* < 0.0001, *ns*. not significant, unpaired Student’s t test (**B**, **C**, **E**, **G**, **H**, **J**). Mean ± SEM.

