## Supplementary figures and images for "ARIH1 deficiency impairs spatial learning and memory via GIRK2 upregulation in hippocampal CaMKII-expressing neurons in mice"

### Supplemental Figure 1

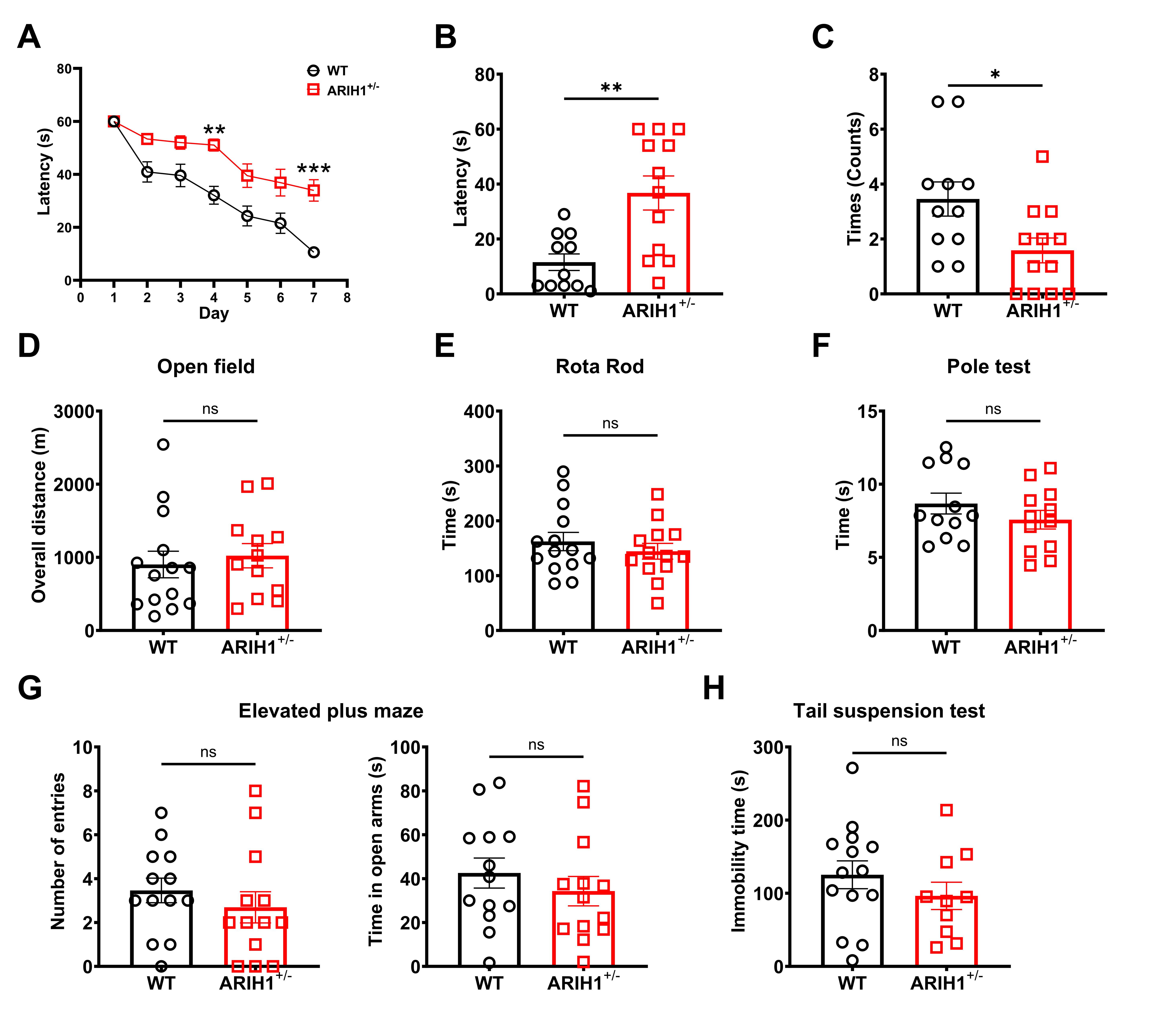

### Supplemental Figure 2

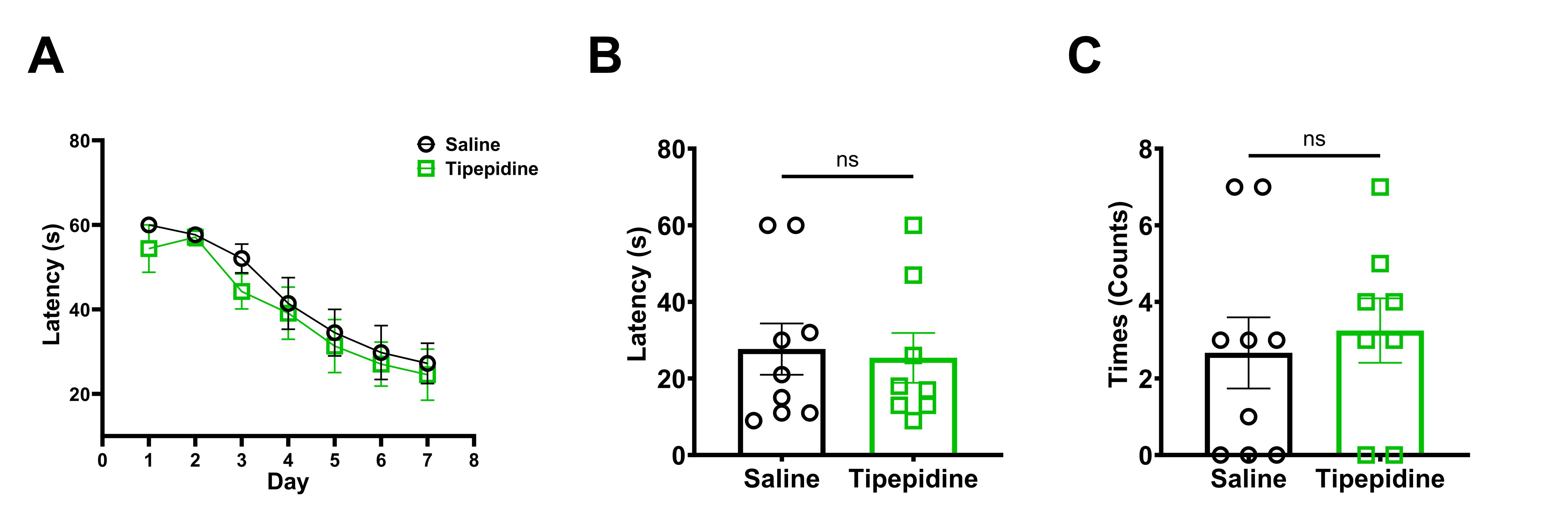

### Supplemental Figure 3

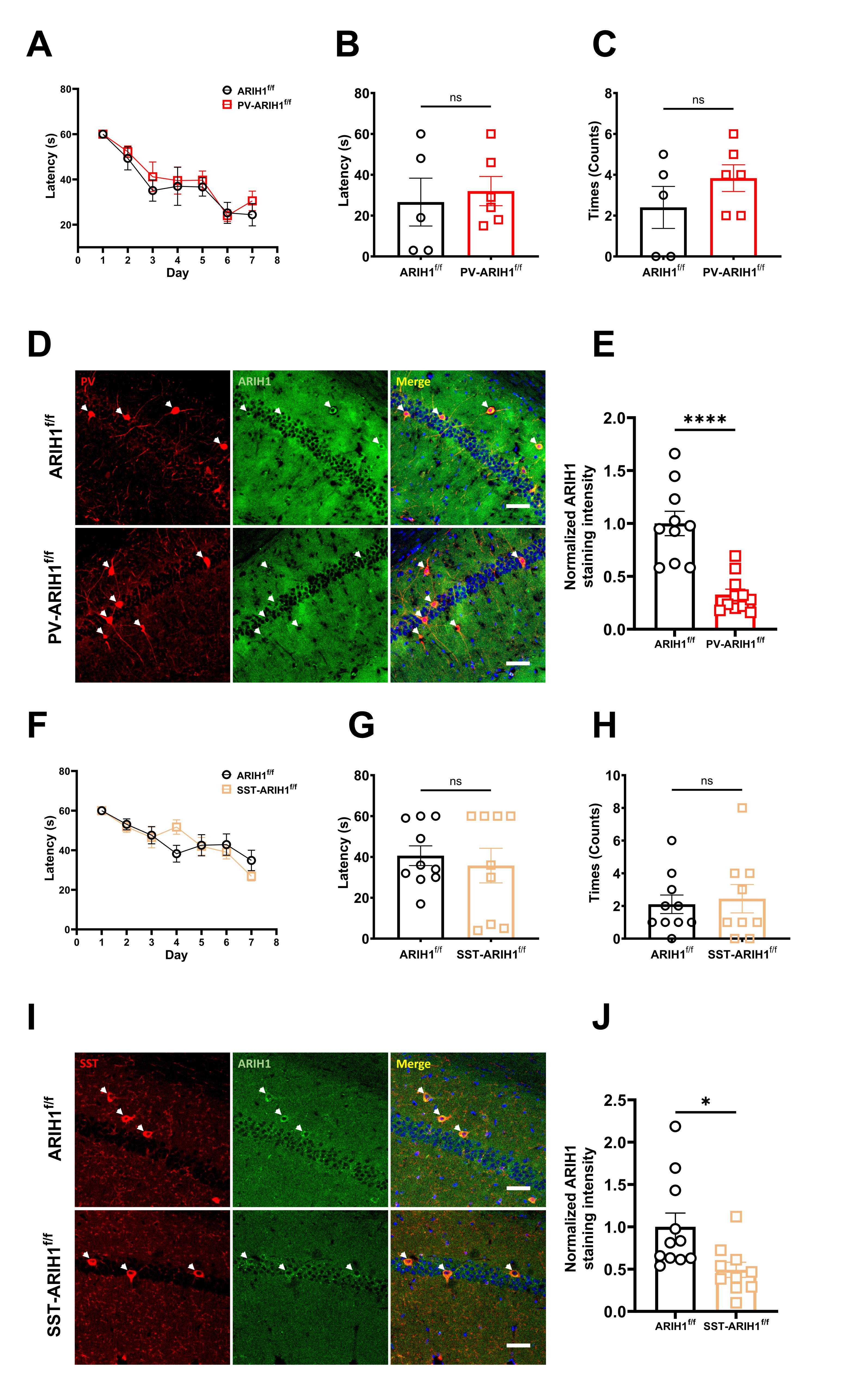
